# Maturation of the glymphatic system confers innate resistance of the brain to Zika virus infection

**DOI:** 10.1101/2025.10.26.684625

**Authors:** Jhulimar Guilherme Doerl, Emanuelly Karine Campos Chaves, Vivian Antonielly Becker dos Santos, Bruna Lorena de Melo Marcelino, Diego Marques Coelho, Leo Morita Miyakoshi, Josélio Maria Galvão de Araújo, Selma Maria Bezerra Jerônimo, Eduardo Bouth Sequerra

## Abstract

The maturation of the glymphatic system initiates fluid dynamics in the central nervous system (CNS). Here, we demonstrate that the initiating glymphatic function results in a steep reduction in the susceptibility of the CNS to Zika virus (ZIKV) infection. In mice, ZIKV injection before glymphatic system maturation leads to widespread infection, vascular deformation, and disruption of Aqp4^+^ astrocytic networks. After the glymphatic system formation, the same challenge fails to cause disease or persistent infection. Pharmacological inhibition of vasopressin receptor 1a, which reduces perivascular fluid dynamics, reopens the window of susceptibility to ZIKV-related disease. These findings reveal the glymphatic system as an innate protective barrier that limits viral spread in the brain parenchyma and suggest a possible explanation for why late-gestation human fetuses and newborns are less susceptible to ZIKV. Moreover, these data indicate that children with ZIKV-caused malformations can have continuous brain inflammation due to a defective glymphatic system.

## Introduction

Right after the introduction of the Zika virus (ZIKV) in the naïve Brazilian population in 2015, multiple human embryos were infected by vertical transmission. Thousands of them developed central nervous system (CNS) malformations, collectively referred to as congenital ZIKV syndrome (CZVS) ^1–4^. The limited information available about the susceptibility window for ZIKV-caused CNS malformations comes from population studies that identified the gestational week in which pregnant women tested positive or the week in which mothers reported experiencing symptoms ^5,7^. These studies reveal that mothers of patients with CZVS were infected in different gestational weeks, covering the whole extent of pregnancy. However, the susceptibility to ZIKV-derived CNS malformations diminished as neural development progressed. The probability of developing neurological and ocular malformations was lower in the third trimester ^6,8^. Moreover, the reports of postnatal ZIKV infections causing neurological consequences are still sparse ^9^.

In adults, a crucial mechanism for managing brain viral infections involves the transport of parenchymal interstitial fluid into the meningeal lymphatic vessels. For instance, both the ligation of the lymphatic vessels to the cervical lymph nodes and the photo ablation of meningeal lymphatic vessels lead to faster death by the Japanese encephalitis virus. In contrast, expanding meningeal lymphatic vessels reduced the number of deaths ^10^. Viruses and other particles in the brain are transported to the meninges via the glymphatic system ^11^. In the glymphatic system, the cerebrospinal fluid (CSF) in the meninges permeates into perivascular spaces formed by astrocyte feet and blood arteries. From the perivascular space, astrocytes transport the fluid to the brain parenchyma, where it mixes with the interstitial fluid. The interstitial fluid is again transported by astrocytes into the perivascular space of blood veins, then reaches the meninges and the lymphatic system ^12^. Here, we hypothesize that the onset of glymphatic function represents a step at which the brain abruptly decreases its susceptibility to ZIKV infection. Since human cortical astrocytes begin to differentiate by the end of the second trimester ^13^, the third trimester is likely crucial for the initiation of glymphatic system function.

In mice, the glymphatic system begins to function postnatally, progressing from the caudal to the anterior regions and from ventral to dorsal ^14^. It is complete by postnatal day 14 (P14), when astrocytes are in place, with their perivascular feet differentiated and expressing Aqp4 ^14^. Astrocytes use Aqp4 to transport water between the perivascular space and the parenchyma ^15^. Coincidentally, the susceptibility of mice to developing disease or dying after intraperitoneal ZIKV injection diminishes between P7 and P14 ^16^.

Here, we show that ZIKV brain injection in the first postnatal week disrupts the glymphatic system formation. In contrast, ZIKV intracerebral injection in the third postnatal week has minimal consequences for the glymphatic system function and the animal’s health. Moreover, we demonstrate that pharmacological inhibition of the glymphatic system, by blocking the V1aR and reducing arterial contractility, in P14 mice reopens the susceptibility window for ZIKV-induced brain disease and the glymphatic system function defect.

## Results

### Mice display age-dependent susceptibility to disease caused by ZIKV brain injection

Mice show decreased susceptibility to developing disease and dying after intraperitoneal injection of ZIKV when they reach postnatal day 14 (P14) ^16^. However, ZIKV replicates in the adult mouse brain and causes weight loss ^17^. Therefore, we investigated whether the brains of C57BL/6 mice exhibit age-dependent susceptibility to ZIKV through direct intracerebral injections. We injected P1 and P21 mice, representing animals with and without a functional glymphatic system ^14^ (Figure 1A). After 13 days, we injected the tracer FITC-Dextran into the cisterna magna (Figure 1A) of ketamine and xylazine-anesthetized animals. We interrupted the experiment 30 minutes later by decapitation and immersing the brain in 4% PFA.

**Figure 1.**
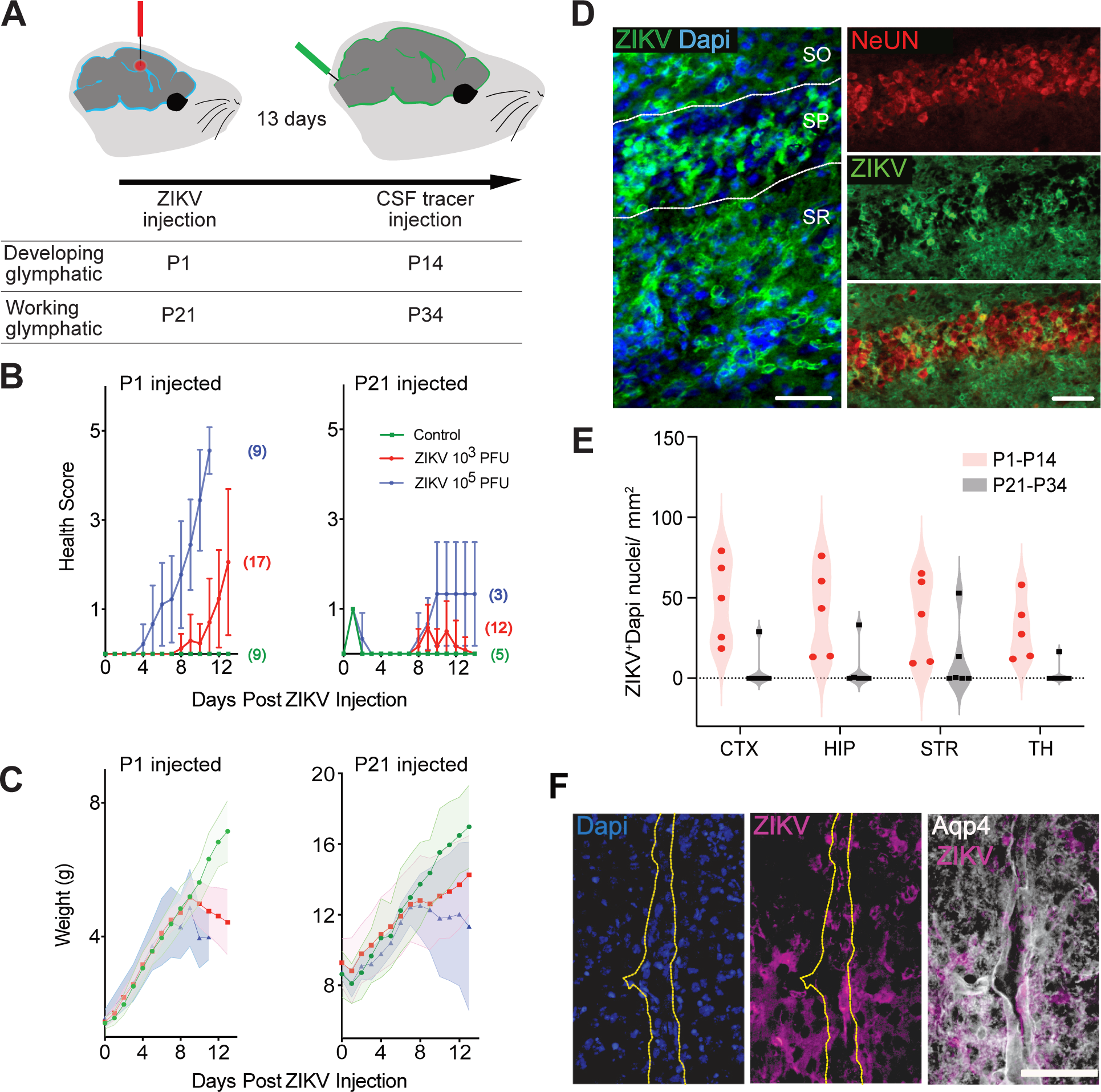
Age-dependent susceptibility to disease caused by ZIKV injection in the brain. **A.** Timeline showing the two experimental groups representing animals before the glymphatic system has developed (P1) and after the glymphatic system is functional (P21). The animals were followed for 13 days, until the experiment was performed to test glymphatic system activity. **B.** Graphs showing the development of disease signs during post-infection days (mean ± SE). The number of animals is represented in parentheses in the color of each group. **C.** Graphs show the weight gain during development (mean ± SE, colored area). Note that P1 animals injected with 10^5^ PFU died before day 13. **D.** Photomicrographs of the CA1 region of the hippocampus of a ZIKV-infected P14 animal. Note that ZIKV-infected cells are spread throughout all the layers. SO. Stratum Oriens, SP. Stratum Pyramidale, SR. Stratum Radiatum. Some of the infected cells are neurons in the SP (NeUN+), but multiple infected cells are NeUN-. **E.** Graph showing the variability in the density of infected cells between P14-infected animals and brain regions, as well as the low frequency of infected animals in the P34 group. **F.** Photomicrograph of a blood vessel region in the cerebral cortex (V1) where astrocyte end feet express Aqp4 (white, yellow dashed lines) in close association with ZIKV-infected cells (purple). Scale bars 50 µm.

We injected two doses of ZIKV, 10^3^ and 10^5^ plaque-forming units (PFU), into the right lateral ventricle. 10^5^ PFU-injected P1 animals began developing disease symptoms by 4 days post-ZIKV infection, and all nine animals died at the 11^th^ day post-infection (Figure 1B). They started losing weight just one day before dying (Figure 1C). On the other hand, 10^5^ PFU-injected P21 animals started developing symptoms around the 8^th^ day post-infection and did not die during the 13-day experiment (Figure 1B). Since ZIKV P1-injected animals did not reach P14, we did not analyze the effect of ZIKV on the glymphatic system after 10^5^ PFU injections.

10^3^ PFU-injected P1 animals started to develop disease signs at eight days post-infection and to lose weight two days before the experiment, while P21-injected animals developed milder symptoms and did not lose weight (Figure 1B and C). Moreover, ZIKV-injected P1 animals began to develop postnatal reflexes, such as decubitus recovery and bar holding, but ceased displaying these reflexes by 13 days post-infection, with almost none of the tested reflexes remaining (Supplementary Figure 1). Therefore, these data indicate an age-dependent susceptibility window for ZIKV-induced disease, and the associated symptoms are likely related to the brain infection.

To determine whether injecting ZIKV at these two ages results in different infection levels, we performed immunohistochemical detection of the ZIKV envelope protein at 13 days post-infection using the 4G2 antibody (Figure 1D). While all the P1-injected animals had varying densities of ZIKV-infected cells in all different brain regions, only one P21-injected animal had ZIKV^+^ cells spread through the four areas analyzed, and another one only in the striatum (Figure 1E). Therefore, P21 animals are less likely to develop a persistent ZIKV brain infection after 13 days, suggesting an internal mechanism for diminishing susceptibility.

ZIKV-infected cells were found in both the brain parenchyma and near Aqp4+ astrocytic feet. This proximity suggests that some infected cells were situated within or near the perivascular space, which is formed between astrocytes and the endothelium of blood vessels (Figure 1F). Therefore, we decided to next test the function of the glymphatic system in animals injected with 10^3^ ZIKV PFU at P1 and P21.

### ZIKV infection disrupts the glymphatic system development but does not deform it after its maturation

The peak of prosencephalic astrocyte generation in mice occurs during the first postnatal week, through the transformation of part of the radial glial cells in the subventricular zone that translocate into the parenchyma ^18–21^. Since ZIKV preferentially infects astrocytes ^22–25^, we tested whether injecting it into the brain disrupts development and/or damages the mature glymphatic system. After 13 days of infecting P1 or P21 mice through the injection of 10^3^ PFU of ZIKV into the right lateral ventricle, we injected FITC-Dextran into the fourth ventricle via the cisterna magna (Figure 1A). We kept the animals alive for 30 minutes and fixed their brains by immersion in 4% PFA overnight. P1-infected animals showed reduced CSF transport into the parenchyma of different brain regions (Figure 2A and C; multiple Mann-Whitney tests, FDR for p-adjustment after multiple comparisons, hippocampus p= 0.000006, cerebral cortex p= 0.00002, dorsal olfactory bulb p= 0.00008, ventral olfactory bulb p= 0.000009, thalamus p= 0.0009, midbrain p= 0.0001, cerebellum p= 0.19, medulla p= 0.0002, striatum p= 0.03, and corpus callosum p= 0.0004), while P21-infected animals showed the same amount of FITC in the parenchyma as controls (Figure 2B and D; multiple Mann-Whitney tests).

**Figure 2.**
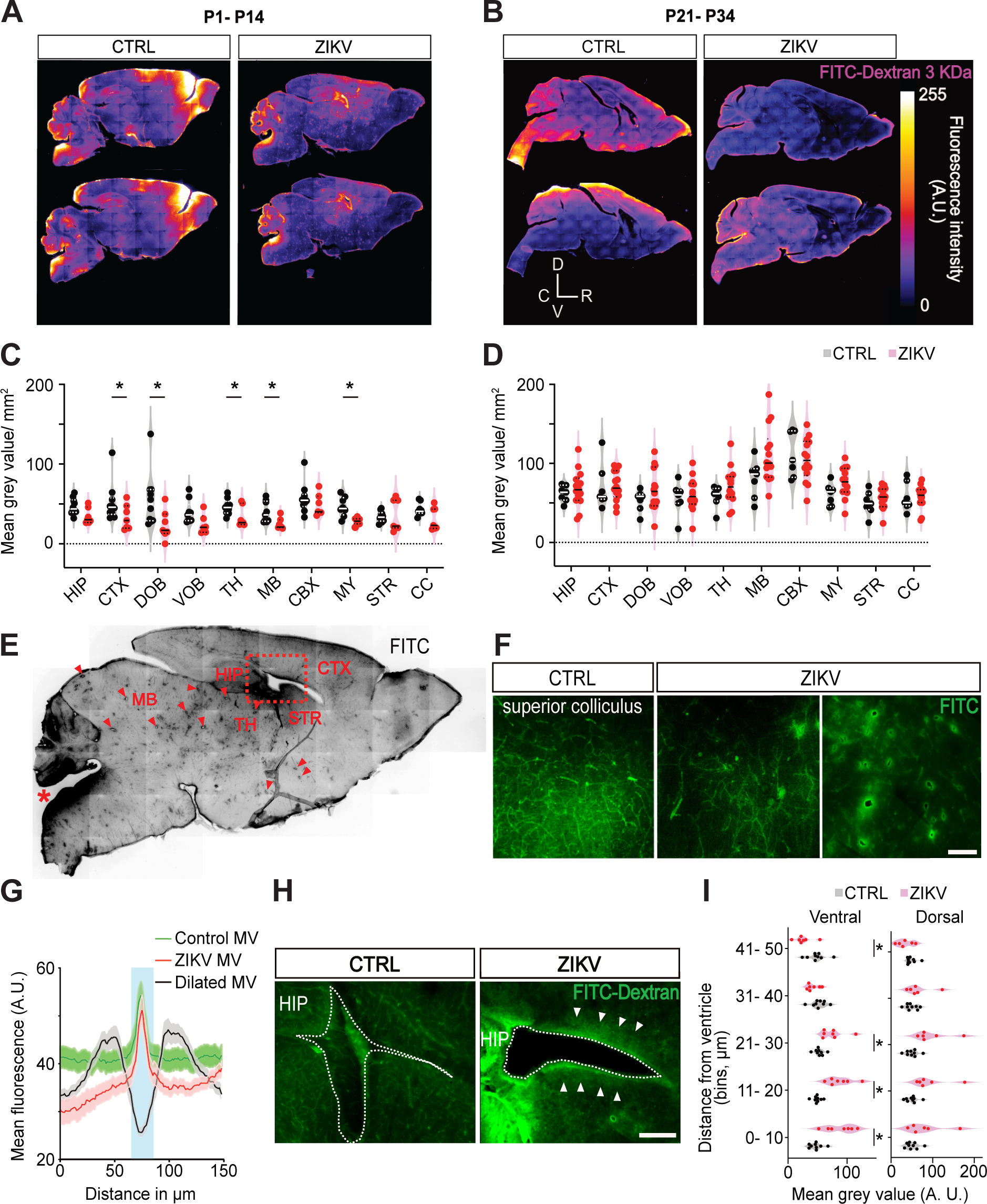
ZIKV disrupts the development but does not deform the glymphatic system. **A.** Representative reconstructions of sagittal slices from P1 saline-injected (CTRL) or animals injected with 10^3^ PFU of Zika virus (ZIKV). Colors indicate the intensity of FITC-derived fluorescence after injection in the cisterna magna at P14. **B.** Representative slices from controls and ZIKV-injected animals in P21 that were later injected with FITC-Dextran in P34. Wind rose indicates dorsal (D), ventral (V), rostral (R), and caudal (C). **C.** ZIKV-injected P1 animals display lower transport of FITC to several brain regions, while those animals injected in P21 display a similar transport of FITC to the brain parenchyma as controls (**D**). Asterisks in the graphs indicate p ≤ 0.05, multiple Mann-Whitney tests, and FDR for p-adjustment after multiple comparisons. **E.** Some P1 ZIKV-injected animals have FITC-stained ellipsoid bodies in the tissue through which the CSF diffuses to the parenchyma. These were located in the midbrain, thalamus, and basal ganglia (arrowheads). The red asterisk indicates the FITC injection site. **F.** Higher magnification of the superior colliculus of one control animal (left) and two P1 ZIKV-injected animals with different levels of modification of FITC transport through the perivascular spaces of microvessels, including the ellipsoid bodies (center, right), scale bar 150 µm. **G.** Plot of the mean fluorescence around micro and dilated microvessels (MV, centered at 75 µm) showing mean ± SE. H. Photomicrograph of the region indicated with the red, dashed rectangle in E. Picture shows the difference in fluorescence around the lateral ventricle (arrowheads), scale bar 500 µm. **I.** Graphs show the FITC diffusion to the ventral ventricular wall (adjacent to the striatum) but not to the dorsal ventricular wall (adjacent to the cerebral cortex). Asterisks in the graph indicate p ≤ 0.05, multiple t tests and FDR for p-adjustment after multiple comparisons. HIP. hippocampus, CTX. cerebral cortex, DOB. dorsal olfactory bulb, VOB. ventral olfactory bulb, TH. thalamus, MB. midbrain, CBX. cerebellum, MY. Medulla oblongata, STR. striatum, CC. corpus callosum.

Although we detected a general reduction in CSF transport to the brain parenchyma in the P1 ZIKV-infected animals, we identified two distinct abnormal FITC signal sources that may indicate the breakdown of barriers. In one of them, we observed ellipsoid spaces that are absent in controls or in P21-infected animals and serve as entry points for FITC (Figure 2E and F; present in six of the seven infected animals). These spaces are mainly located in the midbrain, thalamus, and striatum (Figure 2E; density of 12.16 ± 6.67 SD per mm² in the midbrain, n=6) and have an average perimeter of 124.45 ± 38.86 SD μm. Compared to control microvessels or microvessels in the ZIKV-infected parenchyma, the ellipsoid spaces exhibit a much longer diffusion of FITC into the parenchyma (Figure 2G; 33.15 ± 6.12 SD μm, n=10). The second region with a higher FITC signal we observed was the ventricular region in P1-infected animals (Figure 2H). Using a grid with five 10 μm bins from the ventricular surface into the parenchyma (100 μm wide), we found that FITC enters more from the lateral ventricle into the first 30 μm of the infected striatum than in controls, with an inversion of this relationship occurring at the 40-50 μm bin (Figure 2I; multiple t-tests, FDR for p-adjustment after multiple comparisons, bin 0-10 μm p=0.003, bin 10-20 μm p=0.002, bin 20-30 μm p=0.02, bin 30-40 μm p=0.10, and bin 40-50 μm p=0.01). There is no effect of ZIKV on the permeability of the ventricular wall adjacent to the dorsal subventricular zone (Figure 2I). Therefore, although CSF transport into the parenchyma is reduced in P1-infected animals, multiple points of pathological communication exist between the CSF and interstitial fluid, such as the wall of the striatal subventricular zone and the ellipsoid spaces in the parenchyma, which we describe to be dilated blood vessels (see below).

### ZIKV infection during the first postnatal week disrupts the network of Aqp4^+^ perivascular spaces

Since ZIKV infection during development inhibits glymphatic function at P14, we aimed to describe its effects on astrocytes and their feet near blood vessels, which are essential components of the glymphatic system. The Aqp4 staining in the ZIKV-infected brains showed varying degrees of alteration in its spatial expression pattern compared to controls. While some animals showed mild or no alterations, others exhibited a loss of resolution in Aqp4 expression near blood vessels, and some displayed large, low-density Aqp4 signal areas (Figure 3A). Quantification of the Aqp4 polarity index (fluorescence peak value over the blood vessel divided by the mean fluorescence in the surrounding parenchyma ^26,27^) revealed a reduction in ZIKV-infected animals (Figure 3 B; t-test, p= 0.047). This reduction in polarity correlated with an increased Aqp4 signal in the surrounding parenchyma (Figures 3C and D). Comparing the expression of CD31, a vascular endothelium marker, with Aqp4 showed that ZIKV-infected animals had a reduced length of blood vessels covered by Aqp4^+^ astrocyte feet in various brain regions (Figure 3E; multiple t tests, FDR for p-adjustment after multiple comparisons: p= 0.001 across regions, including the visual cortex, striatum, nucleus accumbens, and motor cortex).

**Figure 3.**
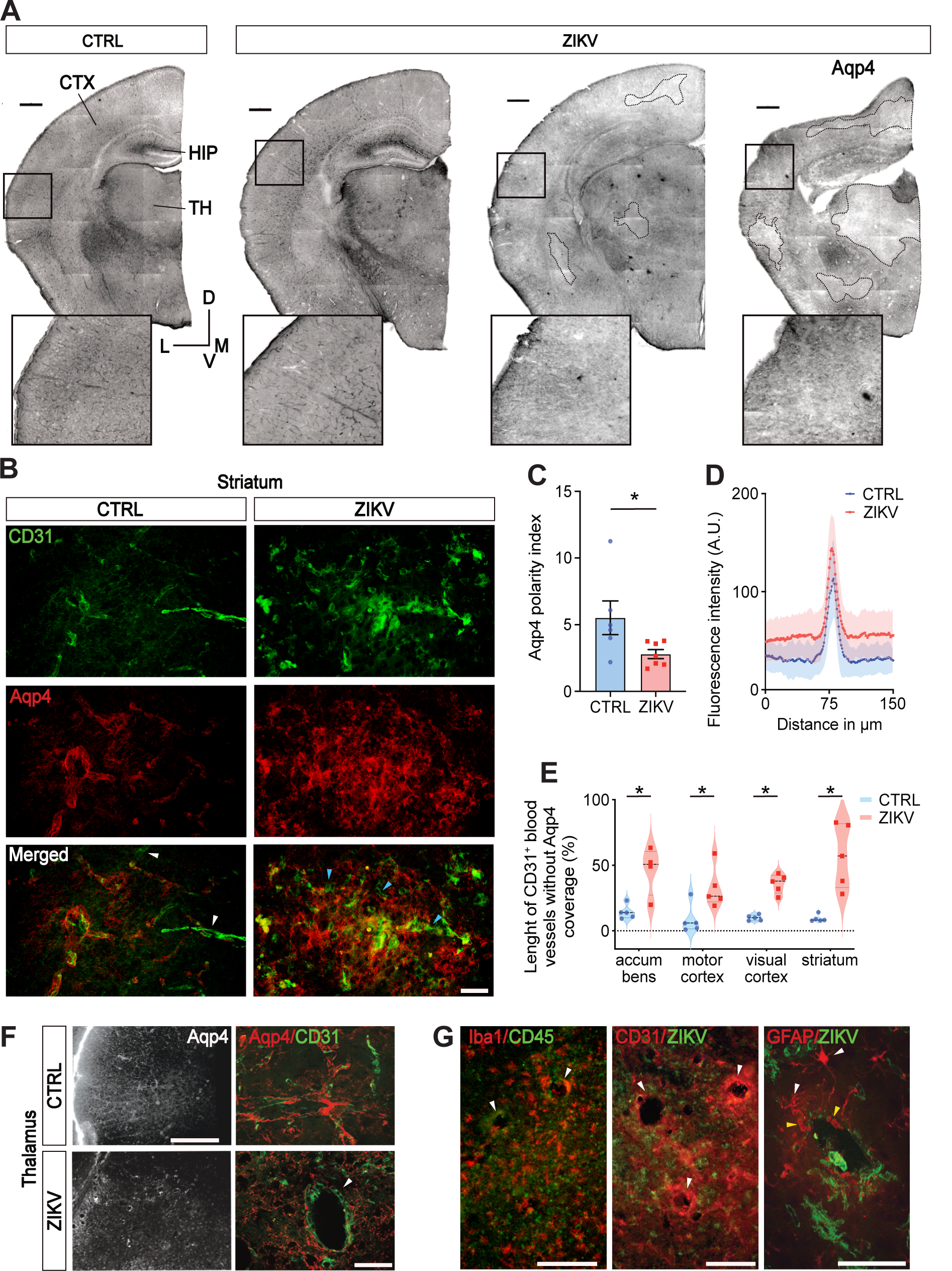
ZIKV disrupts the development of Aqp4+ astrocyte feet around the blood vessels. **A.** Coronal slices showing different levels of alterations in the Aqp4^+^ astrocyte feet around blood vessels in P1 ZIKV-injected mice, analyzed in P14 (three slices on the right), when compared to controls (left slice). Note that the ZIKV-injected animal on the left has maintained Aqp4 polarization towards the blood vessels, while the two on the right had a disorganized expression. The brain on the right was obviously malformed, with cortical thinning and ventricle enlargement. Dashed lines surround Aqp4 low-expression zones. Scale bars = 500 µm. The insets show a higher magnification of the tissue inside the rectangles. **B.** Immunohistochemical labeling of striatal CD31^+^ endothelial blood vessel cells and Aqp4 in astrocytes. White arrowheads show blood vessels devoid of any Aqp4^+^ astrocyte feet in controls, and blue arrowheads show interruptions of Aqp4^+^ feet in some blood vessels of infected animals. Scale bar 50 µm. **C.** Graph shows mean ± SE of the ratio between the peak fluorescence values for Aqp4 over the blood vessels and the median fluorescence in the 150 µm of brain parenchyma surrounding it in the thalamus of different animals (* p≤ 0.05, t-test with Welch’s correction). **D.** The graph shows the superposition of the mean fluorescence ± SE of Aqp4 immunofluorescence of all the micro-vessels (11) analyzed in each animal, shown in C. The center of the vessels was aligned at 75 µm. Note that Aqp4 expression spreads through the parenchyma in ZIKV animals. **E.** Graph showing a violin plot with mean ± SE of the percentage of CD31+ blood vessels that are not covered by Aqp4 expression in each brain area. * p≤ 0.05, multiple t tests with correction for false discovery rate. **F.** Immunohistochemistry for CD31^+^ endothelial cells and Aqp4^+^ astrocytes in the thalamus, showing that the ellipsoid bodies are formed by dilated microvessels that are not present in controls. Scale bar 50 µm. **G.** Immunohistochemical qualitative analysis showing that multiple CD45+ leucocytes infiltrate into the dilated vessels-containing thalamus (left panel, scale bar 250 µm), that the endothelial cells are not positive for the ZIKV envelope protein (middle panel, scale bar 100 µm), nor are the GFAP^+^ astrocytes (right panel, scale bar 100 µm). The white arrowheads point to dilated microvessels, and the yellow arrowheads point to auto-fluorescent red blood cells.

Most of this reduction resulted from interruptions in the Aqp4+ feet network near blood vessels, rather than a complete absence of the Aqp4 signal, as observed in some control vessels (Figure 3D). Thus, ZIKV disrupts the formation of the Aqp4+ astrocytic feet network that constitutes the glymphatic system.

By double-staining CD31 and Aqp4 in the brain regions where we identified the ellipsoid spaces that are permeable to the CSF, we found that these spaces are dilated microvessels (Figure 3F). We observed infiltration of CD45^+^ leukocytes and the presence of red blood cells near these dilated microvessels, indicating increased permeability in these areas (Figure 3G). Additionally, the ZIKV envelope protein was not associated with blood vessel endothelial cells but with cells in the brain parenchyma, and the virus did not infect GFAP^+^ astrocytes at this time point (Figure 3G).

Therefore, infection of the brain during gliogenesis leads to disorganization of the astrocytic network that forms the glymphatic system. At some points, particularly in the midbrain, thalamus, and striatum, the deformation of the perivascular space correlates with vascular endothelium dilation. The perivascular spaces around these dilated vessels are more permeable to CSF.

### The inhibition of the vasopressin receptor 1a increases the brain susceptibility to ZIKV

Since a correlation exists between the severity of brain ZIKV infection and the glymphatic function, we aimed to inhibit brain fluid dynamics to reopen the susceptibility window in P14 mice, immediately after its formation ^14^. A key driver of CSF movement through the glymphatic system is the contraction of meningeal and brain arteries ^28^. Vasopressin receptor 1a (V1Ra) activates contraction in these two types of human arteries *in vitro* ^29^. Moreover, inhibition of V1aR with SR-49059 prevents edema formation after traumatic brain injury ^30^, a process regulated by the influx of CSF from perivascular spaces into the damaged area ^31^. Consequently, we repeated the experiment involving intracerebral injection (i.c.) of 10^3^ ZIKV PFU in P14 mice, followed by CSF tracer injection 13 days later (in P27). We included two groups, in which we added either vehicle or SR-49059 to the same solution that we injected into the lateral ventricle with saline or ZIKV.

P14 ZIKV-injected animals only show some symptoms of the health score in the first two days after ZIKV i.c. injection, but by the 13^th^ day, they are as healthy as controls (Figure 4A). On the other hand, animals that received an intracerebral injection of ZIKV and SR-49059 progressively develop worse symptoms until the end of the experiment (Figure 4A). No animals died. Therefore, inhibiting V1aR reopens the susceptibility window for developing disease caused by ZIKV CNS infection.

**Figure 4.**
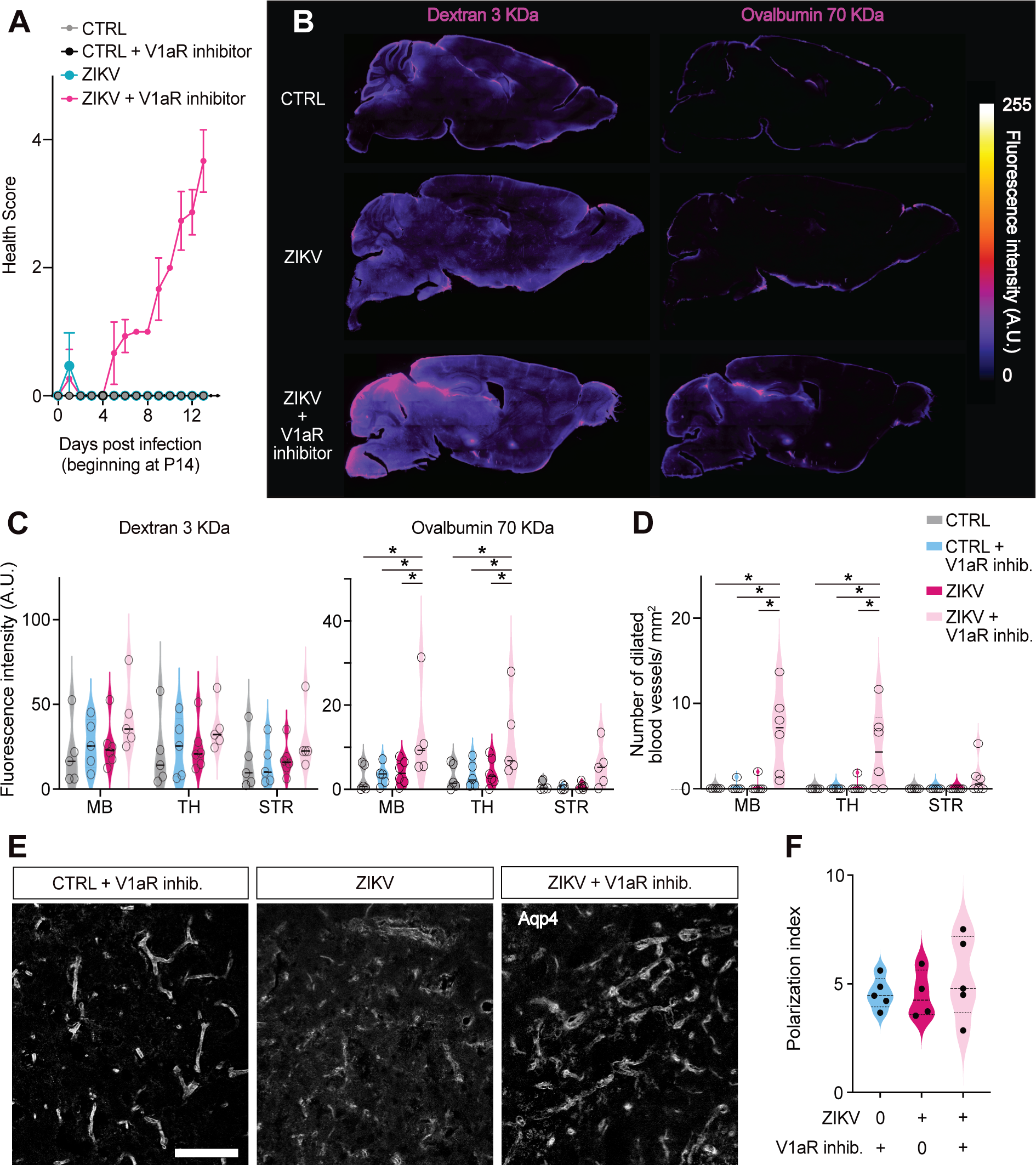
Inhibition of fluid dynamics in the brain reopens the susceptibility window for ZIKV-related disease. **A.** The graph shows the mean ± SE health score of the animals during the 13 days after ZIKV intracerebral injection. **B.** Example reconstructions of brain sagittal slices of a saline-injected (top), a ZIKV-injected (middle), and a ZIKV + SR-49059 (V1aR inhibitor)-injected animal (right). The columns show pictures of the same slice in two channels, Rhodamine-Dextran (left) and ovalbumin-Alexa488 (right). **C.** The two graphs show the mean ± SE fluorescence intensity (mean grey value) of the two tracers across three brain regions (MB - midbrain, TH - thalamus, and STR - striatum) for the four experimental groups. The individual animal means are represented as circles. **D.** The graph shows the number of dilated blood vessels per area in the same three brain regions analyzed in C. * p≤ 0.05, 2-way ANOVA, and Tukey’s multiple comparisons test. **E.** Representative photomicrographs of the Aqp4 immunohistochemistry in the striatum of animals from three experimental groups. Scale bar 50 µm. **F.** The graph shows the mean ± SE Aqp4 polarity index in the striatum for each group. Individual animals are represented as circles.

We then compared the transport of two differently sized CSF-injected tracers in the parenchyma: rhodamine-dextran (3 kDa) and ovalbumin-Alexa488 (70 kDa; Figure 4B). Although dextran transport showed no differences between groups (2-way ANOVA and Tukey’s multiple comparisons test), ovalbumin transport into the parenchyma of the midbrain and thalamus of ZIKV + SR-49059-injected animals increased (Figure 4C; 2-way ANOVA and Tukey’s test, Midbrain: CTRL vs ZIKV + SR-49059 p 0.01; CTRL + SR-49059 vs ZIKV + SR-49059 p 0.02; ZIKV vs ZIKV + SR-49059 p 0.03; Thalamus: CTRL vs ZIKV + SR-49059 p 0.02; CTRL + SR-49059 vs ZIKV + SR-49059 p 0.04; ZIKV vs ZIKV + SR-49059 p 0.04). Additionally, ZIKV + SR-49059-injected animals developed dilated microvessels in the midbrain and thalamus (Figure 4D; 2-way ANOVA and Tukey’s test, Midbrain: CTRL vs ZIKV + SR-49059 p 0.0001; CTRL + SR-49059 vs ZIKV + SR-49059 p 0.0001; ZIKV vs ZIKV + SR-49059 p 0.0001; Thalamus: CTRL vs ZIKV + SR-49059 p 0.005; CTRL + SR-49059 vs ZIKV + SR-49059 p 0.005; ZIKV vs ZIKV + SR-49059 p 0.006).

Because there is greater transport of ovalbumin to the parenchyma in animals injected with ZIKV and SR-49059, we tested whether the network of Aqp4+ astrocyte feet around blood vessels is preserved. We found no difference in the Aqp4 polarity. The groups did not differ in either peak fluorescence or Aqp4 surrounding blood vessels. Analysis of the ZIKV envelope protein by immunohistochemistry revealed that none of the animals in Figure 4E had detectable ZIKV^+^ cells at 13 days post-infection.

### Human astrocytes begin to express Aqp4 in the third trimester, concomitant with the reduction in CZVS cases

Human cortical astrocytes begin to differentiate at the end of the second trimester and are continuously generated until close to birth ^13^. To get a better sense of when the glymphatic system begins to function, we used a publicly available bulk RNA-Seq dataset (Brainspan atlas) from multiple human brain areas across different developmental stages. Our analysis shows that Aqp4 expression increases dramatically between the second and third trimesters and continues to increase after birth (Figure 5A). Therefore, the timing of astrocyte Aqp4 expression onset across multiple brain areas supports our hypothesis that glymphatic system development represents a switch in the brain’s innate immune response to ZIKV.

**Figure 5.**
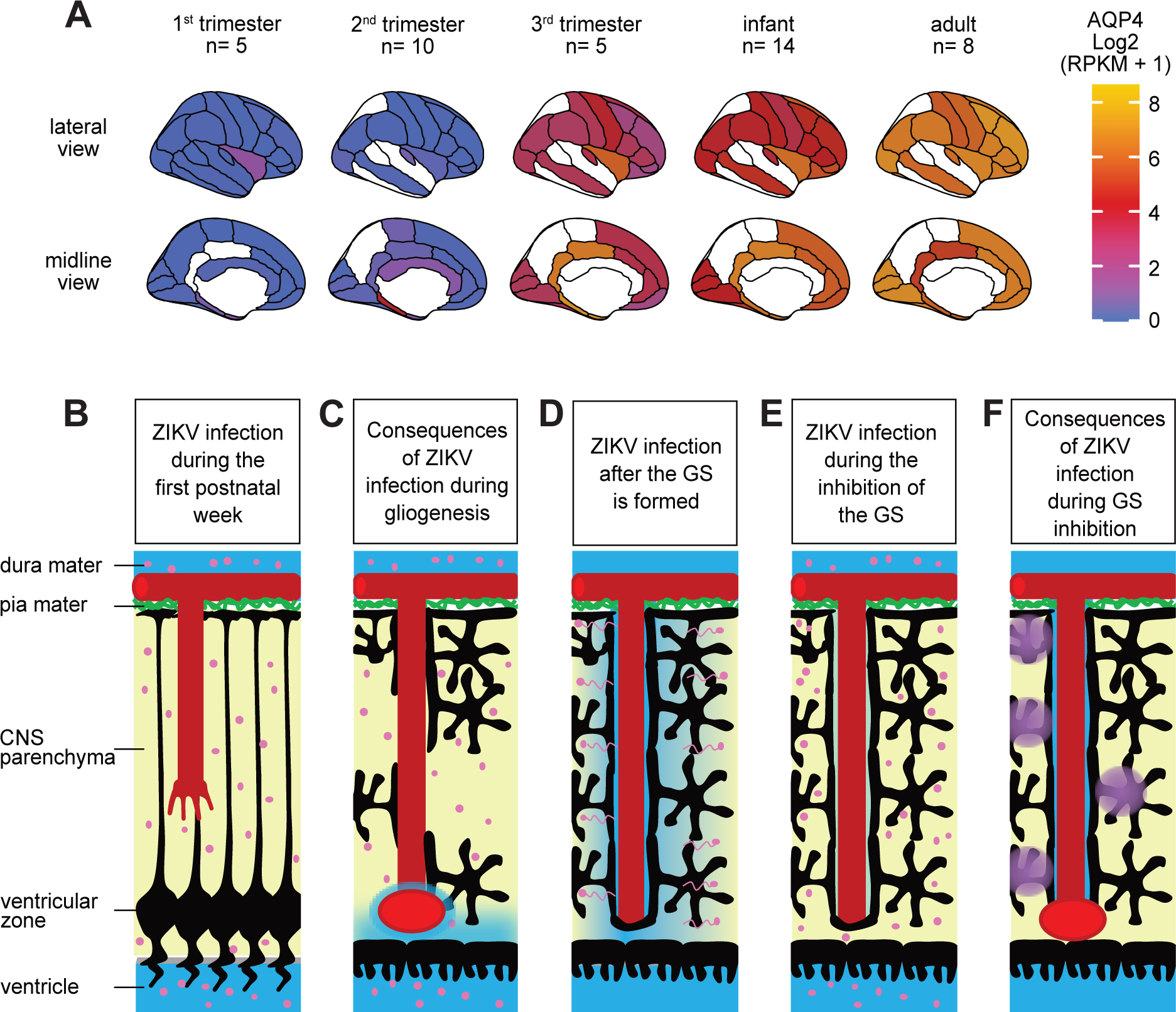
Working hypothesis for the role of the glymphatic system development in the susceptibility of the CNS to ZIKV infection. **A.** The graphical representation of the lateral (top) and medial µm on the right) increases in the third trimester and reaches its highest levels in adults. The RNA-Seq data were obtained from BrainSpan (www.brainspan.org). The color represents the mean expression for the number of individuals represented at each stage. **B.** Graphical representation of the brain parenchyma before gliogenesis, as seen in P1 animals. At this stage, the ventricular surface is formed by radial glial cells (black). These cells project long processes towards the pia mater (green), where they form the barrier between the parenchyma and the meninges (glia limitans). The developing blood vessels at this stage (red) lack a perivascular space around their endothelium. ZIKV (pink circles) infection at this stage causes greater damage because extracellular particles have ample time to infect cells without being cleared. **C.** The effect of ZIKV infection during astrocyte (black) formation is a disorganized network of astrocytes, with poor CSF flow into the perivascular spaces and the brain parenchyma. The main spots of CSF entry in the parenchyma are through dilated blood vessels (red ellipse) and the ventricular surface. **D.** The drawing shows a representation of why we believe P21 animals are less susceptible to ZIKV. In animals with a functional glymphatic system, part of the injected ZIKV and ZIKV released during the lytic phase are washed away from the CNS, reducing the likelihood of infecting neural cells. **E.** If the glymphatic system is pharmacologically inhibited, the probability of each ZIKV extracellular particle infecting a cell increases. **F.** Although P14 animals injected with ZIKV and a V1aR inhibitor develop disease, their brain infection does not persist for 13 days. However, this shorter infection leads to blood vessel dilation and an increase in the transport of protein-based tracers (purple), possibly via a mechanism distinct from Aqp4-mediated water flow.

## Discussion

Here, we show for the first time that the development of the glymphatic system influences the susceptibility of the embryonic CNS to a congenital infection. Although the exact timing of glymphatic system development in humans has not yet been described, the histological description of cortical astrocyte development ^13^ and molecular data showing the onset of Aqp4 expression (Figure 5A) suggest that it occurs during the transition from the third trimester to birth. This timing correlates with the observation that prognostic severity decreases when ZIKV infects the CNS at later stages of fetal development ^6,8^. These findings can impact understanding of other congenital infections, as the onset of CSF transport to the parenchyma and the drainage of interstitial fluid are part of the innate immunity of the developing CNS.

Fluid mobility during fetal development is lower than after birth. The folding neural plate faces the amniotic fluid on its apical side. When the neural tube closes, part of this fluid fills the newly formed ventricular system. It is essential to note that the amniotic fluid can be infected with ZIKV ^32^ and may serve as a route for CNS infection at this early stage. CSF production begins after the development of the choroid plexuses, which occurs between E11 and E14 in mice ^33^ and between the 7^th^ and 29^th^ gestational weeks in humans ^34^. The enclosed amniotic fluid undergoes significant changes in protein composition even before the choroid plexus develops ^35^, likely due to transcellular transport in barrier cells ^36^. Therefore, the CSF is produced much before the development of the glymphatic system. One consequence of CSF production without drainage is the accumulation of proteins. The protein concentration in newborn and preterm CSF is higher than in adult CSF ^37^. The concentration of protein in the CSF of different species peaks after birth and falls until stabilizing in adulthood ^38^. However, both the dextran-conjugated and ovalbumin-conjugated fluorophores injected into the CSF only begin to mix with the murine prosencephalic interstitial fluid postnatally ^14^. Therefore, any pathogen reaching the developing CNS before gliogenesis and the formation of the perivascular spaces by astrocytes finds an environment of low fluid dynamics.

During the stages of neurogenesis, the blood vessels in the meninges extend into the brain parenchyma ^39,40^ in parallel to the radial glia fibers ^41^ (Figure 5B). Here, we demonstrate that if ZIKV infects the brain during this period, specifically during gliogenesis, the glymphatic system becomes malformed, resulting in a disorganized astrocyte network that fails to polarize Aqp4 towards the blood vessels and transport CSF from the perivascular spaces into the brain parenchyma (Figure 5C). We observe variability in the level of astrocyte network disorganization, which may parallel the variable phenotypes observed in the population of congenitally exposed children ^42^. Another important observation from ZIKV P1-injected animals is the occurrence of dilated blood vessels, mainly in the midbrain, thalamus, and striatum, where the CSF tracer diffuses into the parenchyma (Figure 5C). Patients with congenital ZIKV syndrome exhibit increased density of lenticulostriate branches of the middle cerebral artery in the basal ganglia and thalamus, known as lenticulostriate vasculopathy ^43–45^. We propose that the dilated microvessels we observe are consistent with this observation in humans. There is intense infiltration of leucocytes and even the presence of red blood cells at these dilation points. It is possible that the blood-brain barrier is also disrupted there, allowing the CSF tracers to enter the bloodstream. This hypothesis could account for the general decrease in tracer signal in P1 ZIKV-injected animals but requires further investigation.

Our hypothesis is that a significant portion of the ZIKV particles injected into P21 animals is washed out during glymphatic drainage (Figure 5D), a process that repeats during each lytic phase. The inhibition of vascular contractility and the consequent reduction in CSF flow in the perivascular spaces (Figure 5E) allow more ZIKV particles to have time to infect cells. Inhibition of the glymphatic system then reopens the susceptibility window of the CNS to ZIKV. There are other factors influencing susceptibility in treated animals, reflected in milder phenotypes compared to those observed in younger pups. We do not observe changes in the transport of the Dextran-based tracer, suggesting that the structural characteristics of the glymphatic vessels, such as tight junctions, are intact. On the other hand, we observed a change in the transport of the protein-based tracer (Ovalbumin; Figure 5F), suggesting that a different transport mechanism may be affected. In animal models of stroke, activation of cellular transcytosis and endocytosis precedes the disorganization of tight junctions in brain endothelial cells ^46,47^. In fact, blood-injected ZIKV reaches the brain parenchyma of immunocompromised mice without disrupting the blood-brain barrier. Moreover, this study demonstrates that ZIKV crosses a monolayer of brain endothelial cells *in vitro* via transcytosis rather than intercellular passage ^48^. The mechanism underlying astrocyte-mediated transport of perivascular space proteins during ZIKV infection needs to be further explored in future experiments.

Our data from P1-injected animals show that ZIKV-infected brains exhibit a dysfunctional glymphatic system 13 days after infection. However, the patients with congenital ZIKV syndrome do not have ZIKV in their brains anymore. Future studies will investigate the function of the glymphatic system after the animal has resolved the viral infection, to assess the persistence of the effects. P1 mice subcutaneously injected with ZIKV resolve the infection within thirty days, with multiple outcomes ^49^. Both the cases of ZIKV-exposed normocephalic children that develop microcephaly postnatally ^6,50^ and patients that show progression of the malformation, including the evolution to hydrocephaly, after ZIKV infection was resolved ^51,52^, suggest that the brain of congenital ZIKV syndrome patients can be persistently inflamed. One reason for the persistence of inflammation can be a malfunctioning glymphatic system and the associated accumulation of molecules such as lactate, Tau, and Amyloid β ^12,53,54^. Therefore, brains with a malformed glymphatic system can prematurely reproduce the chronic inflammation seen in the degenerating aging brain, where the glymphatic function declines ^26^.

In parallel with the investigation into the persistence of the glymphatic system malformation after ZIKV infection resolution, follow-up imaging studies could focus on anatomical markers, such as Virchow’s perivascular spaces, and on the dynamics of tracers in CZVS patients.

Identifying a possible defective glymphatic system and testing therapeutic approaches could mitigate further damage to the CNS. A recent study describes how the mechanical, non-invasive manipulation of the cervical lymphatics in mice improves CSF drainage ^55^ and may indicate future interventions for CZVS patients.

## Material and Methods

### Animals

Here, we used C57BL/6 mice of both sexes at one, 14, and 21 days of age and followed them for 13 days. The mice had free access to their mothers, and the females received food and water *ad libitum*. The animals were housed in a biosafety level 2 room on a 12h light-dark cycle.

The animals were weighed daily and assigned a Health Score of zero to five: zero-animals without clinical signs or symptoms; one-weight loss when compared to the previous day; two-arched/curved back; three-tremors and/ or difficulty moving; four-limb paralysis; five-moribund or dead, taken to euthanasia.

### Postnatal motor reflexes development evaluation

The neonatal reflexes, decubitus recovery, negative geotaxis, placement by vibrissae, bar holding, and palmar grasp were evaluated from the first day post-infection to the 13^th^ day in animals injected with 10^3^ PFU of ZIKV at P1. In decubitus recovery, the animals were placed belly-up, allowing them to return to their natural position with their belly facing the surface. In negative geotaxis, the ability to balance and turn while facing downward was tested; they were required to turn upward at a 45-degree angle. In placement by vibrissae, the right or left vibrissae are stimulated, and the mouse must move its front paws in the direction stimulated while the hind paws are suspended by the experimenter. The bar-holding test assesses balance, proprioception, and aversion to heights. Palmar grasp is a grasping reflex when an object touches one of the paws. Non-realization or partial realization of the reflex was computed as not realized.

### Virus

The ZIKV strain used in this study was isolated in Pernambuco, Brazil, in 2015 (ZIKV PE/ 243, accession number KX197192 ^56^) and kindly donated by the late Dr. Amilcar Tanuri. The virus was replicated, and the number of plaque-forming units was determined in VERO cell cultures. The supernatant from ZIKV-infected VERO cells was aliquoted and stored at -80 °C.

### Intraventricular ZIKV and SR-49059 injections

P1 animals were ice anesthetized. P14 and P21 animals were anesthetized with an i.p. injection of a ketamine/ xylazine (K/X) mixture (150 and 10 mg/ kg). This anesthetic combination was chosen because, under its effect, the glymphatic system is active ^57^. After that, they received an injection of 1 μL of solution containing 10³ or 10⁵ PFU of ZIKV or saline, using a glass pipette and a manual pressure control system (World Precision Instruments, USA).

The V1aR inhibitor, SR-49059, was diluted in ethanol; the resulting stock solution was diluted to 2 μL for injection (1 μg/ μL, 400 nL of ethanol). The total injection volume was 3 μL (1 μL of ZIKV solution and 2 μL of the SR-49059 solution. Control groups for the SR-49059 solution also received an injection containing 400 nL of ethanol in 3 μL. The injections were applied in the right lateral ventricle.

### Intracisternal injection of CSF tracers

Animals were anesthetized with a K/X mixture. They were positioned with their lower jaw close to the chest to better access the cisterna magna. The hair in the area was shaved. The skin was cleaned with 1% iodine and 70% ethanol. An incision was made to open the skin and separate the muscles along the midline. In this protocol, we used a Nanoliter 2010 injection system (World Precision Instruments; USA). The 2 μL solution was injected at a rate of 23 nL/s. In the first set of experiments, P14 and P34 animals were injected with a 2% Dextran-FITC solution (Merck). In the second set, in which we tested inhibition of the V1aR, we injected a mixture of Dextran-rhodamine (2%; Sigma-Aldrich) and Albumin-Alexa488 (2%; Thermo Fisher). After the injection, the glass capillary was left in place in the cisterna magna for an additional 20 minutes. The capillary was then removed, the animals were decapitated, and the brains were immersed in 4% PFA overnight.

### Immunohistochemistry

The brains were sliced using a Thermo Scientific CryoStar NX70 Cryostat and collected either in free-floating or directly on the slide. For immunohistochemical staining, the tissues were washed three times with 0.01 M Phosphate-Buffered Saline (PBS) for 5 minutes. They were then incubated with a blocking agent (1% bovine serum albumin + 0.05% Triton X-100 diluted in PBS) for 1 hour. The excess blocking solution was removed and the primary antibodies were added (Mouse anti-Flavivirus group envelope protein, GeneTex, 1:1000; Rabbit anti-NeuN, Abcam, 1:1000; Rabbit anti-Aqp4, Sigma-Aldrich, 1:1000; Rat anti-CD31, BD Pharmingen, 1:100; Rat anti-CD45, Invitrogen, 1:500; Rabbit anti-Iba1, Wako, 1:1000; Rabbit anti-GFAP, Dako, 1:300) and kept in a humid chamber at 4°C overnight. The sections were washed three times with PBS for 5 minutes each, then incubated with Invitrogen secondary antibodies at 1:500 (Anti-mouse Alexa 488, Anti-rat Alexa 546, and Anti-rabbit Alexa 647) for 2 hours at 37 °C in a humid chamber protected from light. The slides were washed again with PBS and incubated with a nuclear marker (DAPI, Sigma-Aldrich) for 10 minutes.

### Imaging

Images for fluorescent tracer quantification were captured using an Axio Imager M2 microscope (Zeiss) and the Stereoinvestigator software (MBF Bioscience Inc.).

Immunohistochemistry images were captured using the same microscope equipped with the Apotome function (especially when the DAPI channel was required) or on a confocal microscope (LSM 710/Axio Examiner Z1-Zeiss).

### Image analysis

For the quantification of CSF tracer transport/diffusion into the parenchyma, regions of interest (ROI) with 1000 μm^2^ of area were positioned in different regions with the orientation of a brain atlas. The mean fluorescence intensity was measured using the FIJI software ^58^. Three slices were measured for each animal, and the mean of those represents the fluorescence in each brain region of that animal. The same imaging features were used in all animals. To measure the distance of transport/diffusion of the tracer from the perivascular space of microvessels and dilated vessels, we traced a 150 μm-long line perpendicular to and centered on the vessel, and measured the grey value of the pixel below it using FIJI. Eleven microvessels or dilated vessels were measured from three different slices of each animal.

To quantify ZIKV-positive cells, controls were used to establish the background fluorescence intensity. We measured the number of DAPI-labeled nuclei co-localized with ZIKV envelope immunoreactivity.

For quantification of the Aqp4 signal, we traced a 150 μm-long line perpendicular to and centered on the vessel, and measured the grey value of the pixel directly below it using FIJI. Eleven microvessels were measured from three different slices of each animal. Aqp4 polarization index was calculated with the mean intensity of the peak of fluorescence in the middle of the vessel divided by the mean fluorescence of the parenchyma around it ^26,27^. We quantified the percentage length of CD31-positive microvessels covered with Aqp4-positive astrocytic feet using FIJI.

### Statistics

We tested all data sets for normality using the Kolmogorov-Smirnov test. For the test of differences in CSF tracer transport/diffusion or in Aqp4 coverage in CD31 microvessels, we performed either multiple Mann-Whitney or t-tests, depending on the normality test, with FDR for p-adjustment after multiple comparisons. The comparisons between the four groups in the experiment where the V1aR inhibitor was used were performed by 2-way ANOVA and Tukey’s test. We used Prism (GraphPad) to run the tests and produce the graphs.

### RNA Seq data analysis of AQP4 expression in human brains at different ages

Gene expression data were obtained from the BrainSpan: Atlas of the Developing Human Brain project (www.brainspan.org), which contains RNA-seq read counts normalized to reads per kilobase per million mapped reads (RPKM) and averaged to the gene level. The dataset comprises an expression matrix with genes as rows and samples as columns, a metadata table for genes containing identifiers and annotations, and a metadata table for samples describing donor age, brain structure, and other attributes. Data were processed in R (v4.4.1) using the dplyr, tidyr, ggplot2, and ggseg packages ^59^. Ages were ordered chronologically and grouped into five broad developmental stages: first trimester (8–12 post-conception weeks; pcw), second trimester (13–24 pcw), third trimester (25–37 pcw), infant (4 months to 15 years), and adult (18 to 40 years). Brain structure names were harmonized to match the atlas definitions used in ggseg, enabling brain surface or midline visualization. RNA-seq values were transformed as log₂ (RPKM + 1) prior to plotting in order to stabilize variance and improve interpretability of expression patterns.

## Data availability statement

All data supporting the findings are available upon reasonable request to the corresponding author.

## Ethics declarations

All procedures were approved by the Animal Handling Ethics Committee from the Universidade Federal do Rio Grande do Norte (CEUA-UFRN; 393.011/2024).

## Competing interests

The authors declare no competing interests

## Author contributions

J.G.D., E.B.S., S.M.B.J. and L.M.M. designed the experiments. J.G.D., E.K.C.C., B.L.M.M., V.A.B.S., and L.M.M. executed the experiments and analyzed the data. E.B.S. and S.M.B.J. supervised the experiments. J.M.G.A. produced the virus aliquots used in the experiments. D.M.C. performed the bioinformatics analysis. E.B.S. wrote the manuscript, and all authors reviewed and contributed to the text.

## Aknowledgments

We want to thank Dr. Cecilia Hedin-Pereira for the critical comments on the manuscript. We dedicate this work to the loving memory of Mariana Campelo Medeiros, who was the chief of the animal facility during the experiments, and to Dr. Amilcar Tanuri, who was a driving force behind the study of the Zika virus in Brazil. J.G.D. and B.L.M.M. received a PhD fellowship from Coordenação de Aperfeiçoamento de Pessoal de Nível Superior (CAPES). V.A.B.S. received an undergraduate scientific initiation fellowship from Maternidade Escola Januário Cicco, Empresa Brasileira de Serviços Hospitalares. L.M.M. received a postdoctoral fellowship from Conselho Nacional de Desenvolvimento Científico (CNPq-446325/2024-4). S.M.B.J received grants from the National Institute of Science and Technology on Tropical Diseases (INCT-DT; 408294/2024-8) and CNPq (ZIKA-440893/2016-0). E.B.S. received an Early Career Return Grant (CRP/BRA18-05_EC) and a Research Grant (CRP/BRA24-02) from the International Centre for Genetic Engineering and Biotechnology (ICGEB).

---

**Supplementary Figure 1.**
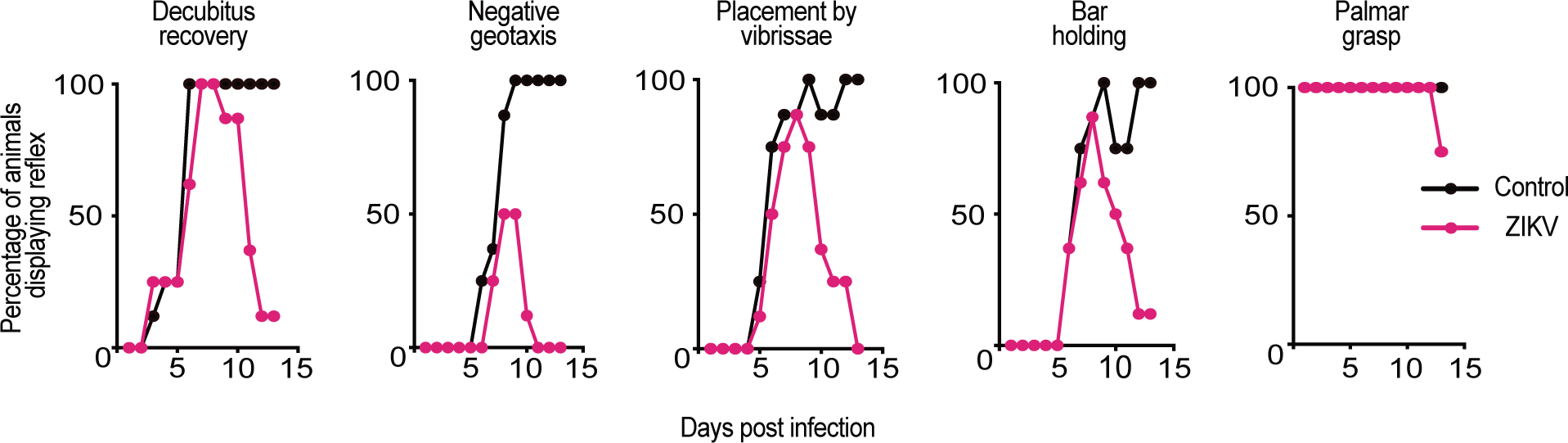
ZIKV infection disrupts the postnatal development of reflexive behaviors. Graphs show the time course of five postnatal reflexes in controls (saline-injected animals; black lines and points) and ZIKV-injected animals (pink lines and points). Data is shown as the percentage of animals in each group that displayed the reflex on each day. Experiments were interrupted at 13 days post-infection for analysis of the glymphatic system. Note that ZIKV-injected animals begin to display the reflexes, but over time, they lose the ability to perform them.

## Notes

### Competing Interest Statement

The authors have declared no competing interest.

## References

1. Aragao, M. de F. V., et al. Clinical features and neuroimaging (CT and MRI) findings in presumed Zika virus related congenital infection and microcephaly: retrospective case series study. BMJ 353, (2016).

2. Aragao, M. F. V. V. et al. Spectrum of Spinal Cord, Spinal Root, and Brain MRI Abnormalities in Congenital Zika Syndrome with and without Arthrogryposis. AJNR Am J Neuroradiol 38, 1045–1053 (2017).

3. França, G. V. A. et al. Congenital Zika virus syndrome in Brazil: a case series of the first 1501 livebirths with complete investigation. The Lancet 388, 891–897 (2016).

4. Moore, C. A. et al. Characterizing the Pattern of Anomalies in Congenital Zika Syndrome for Pediatric Clinicians. JAMA Pediatr 171, 288–295 (2017).

5. Brasil, P. et al. Zika Virus Infection in Pregnant Women in Rio de Janeiro. New England Journal of Medicine 375, 2321–2334 (2016).

6. Nielsen-Saines, K. et al. Delayed childhood neurodevelopment and neurosensory alterations in the second year of life in a prospective cohort of ZIKV-exposed children. Nat Med 25, 1213–1217 (2019).

7. Sequerra, E. B. et al. Association between brain morphology and electrophysiological features in Congenital Zika Virus Syndrome: A cross-sectional, observational study - EClinicalMedicine. https://www.thelancet.com/journals/eclinm/article/PIIS2589-5370(20)30252-2/fulltext?fbclid=IwAR1vzLXpkHp9xbT7TtmOBUKb4AqpoO6nyQ1cijWWXD_vaBjIq6fm3eLPSbM (2020).

8. Hoen, B. et al. Pregnancy Outcomes after ZIKV Infection in French Territories in the Americas. New England Journal of Medicine 378, 985–994 (2018).

9. Ramond, A. et al. Postnatal symptomatic Zika virus infections in children and adolescents: A systematic review. PLoS Negl Trop Dis 14, e0008612 (2020).

10. Li, X. et al. Meningeal lymphatic vessels mediate neurotropic viral drainage from the central nervous system. Nat Neurosci 25, 577–587 (2022).

11. Hablitz, L. M. & Nedergaard, M. The Glymphatic System: A Novel Component of Fundamental Neurobiology. J Neurosci 41, 7698–7711 (2021).

12. Iliff, J. J. et al. A paravascular pathway facilitates CSF flow through the brain parenchyma and the clearance of interstitial solutes, including amyloid β. Sci Transl Med 4, 147ra111 (2012).

13. deAzevedo, L. C., et al. Cortical radial glial cells in human fetuses: depth-correlated transformation into astrocytes. J Neurobiol 55, 288–298 (2003).

14. Munk, A. S. et al. PDGF-B Is Required for Development of the Glymphatic System. Cell Rep 26, 2955–2969.e3 (2019).

15. Mestre, H. et al. Aquaporin-4-dependent glymphatic solute transport in the rodent brain. Elife 7, e40070 (2018).

16. Li, S. et al. Zika Virus Fatally Infects Wild Type Neonatal Mice and Replicates in Central Nervous System. Viruses 10, 49 (2018).

17. Figueiredo, C. P. et al. Zika virus replicates in adult human brain tissue and impairs synapses and memory in mice. Nat Commun 10, 3890 (2019).

18. Suzuki, S. O. & Goldman, J. E. Multiple cell populations in the early postnatal subventricular zone take distinct migratory pathways: a dynamic study of glial and neuronal progenitor migration. J Neurosci 23, 4240–4250 (2003).

19. Alves, J. A. J., Barone, P., Engelender, S., Fróes, M. M. & Menezes, J. R. L. Initial stages of radial glia astrocytic transformation in the early postnatal anterior subventricular zone. J Neurobiol 52, 251–265 (2002).

20. Levison, S. W., Chuang, C., Abramson, B. J. & Goldman, J. E. The migrational patterns and developmental fates of glial precursors in the rat subventricular zone are temporally regulated. Development 119, 611–622 (1993).

21. Luskin, M. B. & McDermott, K. Divergent lineages for oligodendrocytes and astrocytes originating in the neonatal forebrain subventricular zone. Glia 11, 211–226 (1994).

22. Qian, X. et al. Brain-Region-Specific Organoids Using Mini-bioreactors for Modeling ZIKV Exposure. Cell 165, 1238–1254 (2016).

23. Retallack, H. et al. Zika virus cell tropism in the developing human brain and inhibition by azithromycin. Proc. Natl. Acad. Sci. U.S.A. 113, 14408–14413 (2016).

24. van den Pol, A. N., Mao, G., Yang, Y., Ornaghi, S. & Davis, J. N. Zika Virus Targeting in the Developing Brain. J Neurosci 37, 2161–2175 (2017).

25. Watanabe, M. et al. Self-Organized Cerebral Organoids with Human-Specific Features Predict Effective Drugs to Combat Zika Virus Infection. Cell Rep 21, 517–532 (2017).

26. Kress, B. T. et al. Impairment of paravascular clearance pathways in the aging brain. Ann Neurol 76, 845–861 (2014).

27. Lundgaard, I. et al. Beneficial effects of low alcohol exposure, but adverse effects of high alcohol intake on glymphatic function. Sci Rep 8, 2246 (2018).

28. Mestre, H. et al. Flow of cerebrospinal fluid is driven by arterial pulsations and is reduced in hypertension. Nat Commun 9, 4878 (2018).

29. Edvinsson, J. C. A., Grubor, I., Maddahi, A. & Edvinsson, L. Male-female comparison of vasomotor effects of circulating hormones in human intracranial arteries. J Headache Pain 25, 216 (2024).

30. Marmarou, C. R. et al. Selective vasopressin-1a receptor antagonist prevents brain edema, reduces astrocytic cell swelling and GFAP, V1aR and AQP4 expression after focal traumatic brain injury. Brain Research 1581, 89–102 (2014).

31. Mestre, H. et al. Cerebrospinal fluid influx drives acute ischemic tissue swelling. Science 367, eaax7171 (2020).

32. Calvet, G. et al. Detection and sequencing of Zika virus from amniotic fluid of fetuses with microcephaly in Brazil: a case study. The Lancet Infectious Diseases 16, 653–660 (2016).

33. Sturrock, R. R. A morphological study of the development of the mouse choroid plexus. J Anat 129, 777–793 (1979).

34. Bitanihirwe, B. K. Y., Lizano, P. & Woo, T.-U. W. Deconstructing the functional neuroanatomy of the choroid plexus: an ontogenetic perspective for studying neurodevelopmental and neuropsychiatric disorders. Mol Psychiatry 27, 3573–3582 (2022).

35. Chau, K. F. et al. Progressive Differentiation and Instructive Capacities of Amniotic Fluid and Cerebrospinal Fluid Proteomes following Neural Tube Closure. Developmental Cell 35, 789–802 (2015).

36. Parvas, M., Parada, C. & Bueno, D. A blood–CSF barrier function controls embryonic CSF protein composition and homeostasis during early CNS development. Developmental Biology 321, 51–63 (2008).

37. Saunders, N. R. et al. The rights and wrongs of blood-brain barrier permeability studies: a walk through 100 years of history. Front Neurosci 8, 404 (2014).

38. Saunders, N. R. Ontogenetic Development of Brain Barrier Mechanisms. in Physiology and Pharmacology of the Blood-Brain Barrier (ed. Bradbury, M. W. B.) 327–369 (Springer, Berlin, Heidelberg, 1992). doi:10.1007/978-3-642-76894-1_14.

39. Marin-Padilla, M. Early vascularization of the embryonic cerebral cortex: Golgi and electron microscopic studies. J Comp Neurol 241, 237–249 (1985).

40. Marín-Padilla, M. The human brain intracerebral microvascular system: development and structure. Front Neuroanat 6, 38 (2012).

41. Gerhardt, H. et al. Neuropilin-1 is required for endothelial tip cell guidance in the developing central nervous system. Developmental Dynamics 231, 503–509 (2004).

42. Marcelino, B. L. M. et al. Zika virus infection histories in brain development. Dis Model Mech 16, dmm050005 (2023).

43. Mulkey, S. B. et al. Sequential Neuroimaging of the Fetus and Newborn With In Utero Zika Virus Exposure. JAMA Pediatr 173, 52–59 (2019).

44. Nogueira, M. L. et al. Adverse birth outcomes associated with Zika virus exposure during pregnancy in São José do Rio Preto, Brazil. Clin Microbiol Infect 24, 646–652 (2018).

45. Soares de Souza, A., et al. Fetal Infection by Zika Virus in the Third Trimester: Report of 2 Cases. Clin Infect Dis 63, 1622–1625 (2016).

46. Nag, S., Venugopalan, R. & Stewart, D. J. Increased caveolin-1 expression precedes decreased expression of occludin and claudin-5 during blood–brain barrier breakdown. Acta Neuropathol 114, 459–469 (2007).

47. Knowland, D. et al. Stepwise Recruitment of Transcellular and Paracellular Pathways Underlies Blood-Brain Barrier Breakdown in Stroke. Neuron 82, 603–617 (2014).

48. Papa, M. P. et al. Zika Virus Infects, Activates, and Crosses Brain Microvascular Endothelial Cells, without Barrier Disruption. Front. Microbiol. 8, (2017).

49. Ireland, D. D. C. et al. Long-term persistence of infectious Zika virus: Inflammation and behavioral sequela in mice. PLoS Pathog 16, e1008689 (2020).

50. Siqueira Mello, A., et al. Development of Secondary Microcephaly After Delivery: Possible Consequence of Mother-Baby Transmission of Zika Virus in Breast Milk. Am J Case Rep 20, 723–725 (2019).

51. Jucá, E. et al. Hydrocephalus associated to congenital Zika syndrome: does shunting improve clinical features? Childs Nerv Syst 34, 101–106 (2018).

52. van der Linden, V. et al. Association of Severe Hydrocephalus With Congenital Zika Syndrome. JAMA Neurol 76, 203–210 (2019).

53. Iliff, J. J. et al. Impairment of glymphatic pathway function promotes tau pathology after traumatic brain injury. J Neurosci 34, 16180–16193 (2014).

54. Lundgaard, I. et al. Glymphatic clearance controls state-dependent changes in brain lactate concentration. J Cereb Blood Flow Metab 37, 2112–2124 (2017).

55. Jin, H. et al. Increased CSF drainage by non-invasive manipulation of cervical lymphatics. Nature 643, 755–767 (2025).

56. Donald, C. L. et al. Full Genome Sequence and sfRNA Interferon Antagonist Activity of Zika Virus from Recife, Brazil. PLoS Negl Trop Dis 10, e0005048 (2016).

57. Xie, L. et al. Sleep drives metabolite clearance from the adult brain. Science 342, 373–377 (2013).

58. Schindelin, J., et al. Fiji: an open-source platform for biological-image analysis. Nat Methods 9, 676–682 (2012).

59. Mowinckel, A. M. & Vidal-Piñeiro, D. Visualization of Brain Statistics With R Packages ggseg and ggseg3d. Advances in Methods and Practices in Psychological Science 3, 466–483 (2020).

